# Fatty acid metabolic reprogramming promotes *C. elegans* development

**DOI:** 10.1101/2020.12.30.424804

**Authors:** Xuwen Cao, Yusu Xie, Beining Xue, Hanwen Yang, L. Rene Garcia, Liusuo Zhang

**Affiliations:** CAS Key Laboratory of Experimental Marine Biology, Institute of Oceanology, Chinese Academy of Sciences, Qingdao, 266071, China; Laboratory of Marine Biology and Biotechnology, Qingdao National Laboratory for Marine Science and Technology, Qingdao, 266237, China; Center for Ocean Mega-Science, Chinese Academy of Sciences, 7 Nanhai Road, Qingdao, 266071, China; University of Chinese Academy of Sciences, Beijing, 100049, China; Department of Biology, Texas A&M University, College Station, Texas 77843-3258, USA

**Keywords:** acetylcholine signaling, nutrition sensation, developmental regulation, fatty acid metabolism

## Abstract

Acetylcholine signaling has been reported to play essential roles in animal metabolic regulation and disease affected by diets. However, the underlying mechanisms that how diets regulate animal physiology and health are not well understood. Here we found that the acetylcholine receptor gene *eat-2* was expressed in most of the pharyngeal muscles, which is in accordance to our previous report that EAT-2 received synaptic signals not only from pharyngeal MC neurons. The expression of fatty acid synthesis genes was significantly increased in both *eat-2* and *tmc-1* fast-growth mutants on CeMM food environment, compared to the wild-type. Excitingly, dietary fatty acids such as 15-methyl-hexadecanoic acid (C17ISO), palmitic acid (PA, C16:0) and stearic acid (SA, C18:0) supplementation, significantly accelerated wild-type worm development on CeMM, indicating that the fatty acid synthesis reprogramming is an essential strategy for *C. elegans* to regulate its development and growth on CeMM diet. Furthermore, we found that fatty acid elongase gene *elo-6* knock-out significantly attenuated *eat-2* mutant’ fast growth, while overexpression of *elo-6* could rescue the *eat-2; elo-6* double mutant’ slow development, which suggested that *elo-6* played a major role in the above metabolic remodeling. Taken together, our report indicates that diets regulate neuromuscular circuit and modulate *C. elegans* development via fatty acid metabolic reprogramming. As most of the key genes and metabolites found in this study are conserved in both invertebrate and vertebrate animals, we believed that our results might provide essential clues to the molecular mechanisms underlying interactions among animal nutrition sensation, metabolism reprogramming and developmental regulation.

**Significance Statement:** Diets and nutritional composition affect animal development and human health, however the underlying mechanisms remain elusive. We demonstrate that the acetylcholine receptor gene *eat-2* is expressed in most of pharyngeal muscles, and the expression of fatty acid synthesis genes is significantly increased in both *eat-2* and *tmc-1* fast-growth mutants on the synthetic chemical defined CeMM food environment. Dietary supplementation of several fatty acids significantly speed up animal development. Furthermore, we demonstrate that the fatty acid elongase gene *elo-6* knock-out attenuates *eat-2* mutant’ fast growth, and overexpression of wild-type *elo-6* promotes the *eat-2; elo-6* double mutant’ slow development. Our findings describe that acetylcholine signaling coordinate nutrition sensation and developmental regulation through fatty acid metabolic remodeling.

## Introduction

Diet and nutrition affect the health and diseases of animals and humans (1, 2). However, the exact mechanisms of how dietary intake and nutrition composition regulate animal growth, development, reproduction and aging remain elusive. The acetylcholine signaling has been shown to play a critical role in the regulation of diet and energy balance (3, 4). It has been reported that impairment of cholinergic neurons in mouse basal forebrain increases food intake and lead to obesity, whereas enhanced cholinergic signaling reduces food consumption (4). The known appetite suppressant nicotine decreases food intake through activation of nicotinic acetylcholine receptors (5). Additionally, a study in rodents indicates that the addictive properties of nicotine and its diabetes-promoting actions also depend on acetylcholine signaling (6). However, the physiological roles of acetylcholine signaling in diets induced life processes changes, are largely unknown.

Free-living bacterial-feeding *C. elegans* is an important model organism to study how diet and nutrition regulate animal health and life history traits (7–12). The complete connectome makes *C. elegans* an essential model for studying relationships between neuro-regulation and diet (13, 14). *C. elegans* maintenance medium (CeMM)is an axenic chemically defined food source that excludes variables associated with bacterial metabolism (15, 16). Although CeMM has sufficient nutrients, wild-type worms grow slowly and males mate poorly (16, 17). Our previous study has shown that either human and mouse deafness homolog gene *tmc-1* (expressed in cholinergic MC neurons and body wall muscles) or its downstream acetylcholine receptor *eat-2* gene (expressed in pharyngeal muscles) attenuates the development of *C. elegans* worms on CeMM (18). However, the underlying mechanism of how *eat-2* regulates *C. elegans* development is largely unexplored.

In this report, we found that *eat-2* was expressed in multiple pharyngeal muscles, compared to the few expressed puncta near the junction of pharyngeal muscles pm4 and pm5 as previously reported (19). Surprisingly, we found that the fatty acid metabolism of both *eat-2* and *tmc-1* mutants were reprogrammed to accelerate *C. elegans* development. Subsequent fatty acids dietary supplementation, fatty acid elongase genome-editing and rescue experiments indicated that diets regulated neuromuscular circuits and modulated *C. elegans* development via fatty acid metabolic reprogramming. As most of the key genes and metabolic pathways mentioned above are conserved in humans, we believed that our results might provide important clinical clues to human health and diseases.

## Results

### Acetylcholine receptor gene *eat-2* slows down growth on CeMM

*eat-2* is a nicotinic acetylcholine receptor gene, and expressed in pharyngeal muscle of *C. elegans* (19). Mutations in *eat-2* cause *C. elegans* feeding behavior defect (20), and the *eat-2* mutants are widely used as a genetic model for dietary restriction and life span studies (21). We previously found that the *eat-2* mutant grows much faster in the axenic CeMM diet, compared to wild-type N2 worms (18). To study the underlying mechanisms, we prepared CeMM plates according to Zhang *et al.* (18). Since a few chemicals were commercially unavailable in China, nucleic acid substituents Cytidine 3’(2’)-phosphoric acid and Guanosine 2’- & 3’-monophosphate mixed isomers sodium salt were replaced by analogues Cytidine 5’-monophosphate and Guanosine 5’-monophosphate disodium salt hydrate, respectively. In addition, some chemicals were from different manufacturers from those in Zhang *et al.* (18)(SI Appendix, Table S2). In particular, L-lysine monohydrochloride was replaced by DL-lysine monohydrochloride by an unexpected error. All the above chemical substitutions led to slower development than previously reported (18), for both N2 and *eat-2* mutant. In our hands, less than 3% of N2 developed into adults after 9 days post hatching (Fig. 1A) and about 47% of *eat-2* mutants developed into adults by day 6 (Fig. 1B). While 13% of N2 developed into adults on day 8 and 97% of *eat-2* developed into adults on day 5, as reported previously (18). Nevertheless, *eat-2* mutant showed much faster growth than wild-type N2 worms (Fig. 1A and B).

**Fig. 1.**
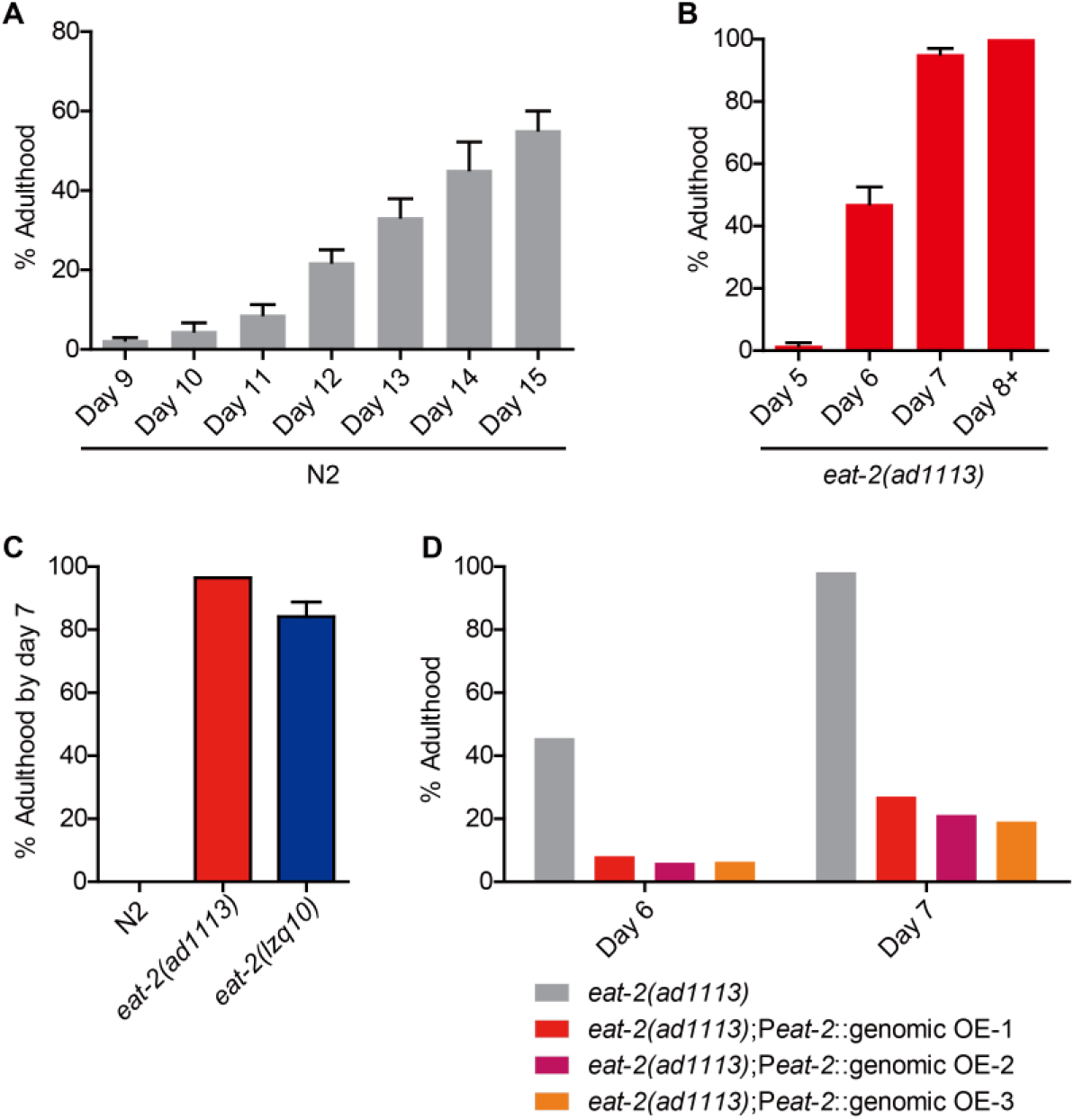
*eat-2* attenuates *C. elegans* development on CeMM diets. (A) Growth rate of wild-type N2 on CeMM media. Three trials, n = 54, 58 and 57 worms, respectively. Growth rate of *eat-2(ad1113)* mutant on CeMM media. Three trials, n = 63, 62 and 62, respectively. (C) The growth rate of *eat-2(lzq10)* on CeMM was similar to *eat-2(ad1113)*. N2, one trial, n = 116; *eat-2(ad1113)*, one trial, n = 57; *eat-2(lzq10)*, three trials, n = 53, 56 and 57, respectively. (D) Transgenic rescue of *eat-2(ad1113)* mutant on CeMM. One trial for four groups, *eat-2(ad1113)*, n = 57; three *eat-2(ad1113)*; P *eat-2*::genomic OE (overexpression) lines, n = 37, 33 and 47, respectively. The value represents the percentage of adulthood per day. Data shown are mean ± s.e.m.

Using CRISPR/Cas9-mediated genome editing, we generated a strain *eat-2(lzq10)* encoding the same R324W mutation as *eat-2(ad1113)*. We recreated this allele to validate that the CeMM growth phenotype was due to the *eat-2* defect, and not due to an unknown linked mutation. Similar to *eat-2 (ad1113)* mutant, the *eat-2(lzq10)* mutant grows much faster compared to wild-type N2 worms on CeMM (Fig. 1C). In addition, we observed that overexpression of wild-type *eat-2* genomic DNA driven by a 4 kb *eat-2* promoter suppressed the fast growth phenotype of *eat-2(ad1113)* mutant (Fig. 1D). Collectively, we confirmed that *eat-2* is a key gene regulating *C. elegans* development in chemically define synthetic food CeMM.

### *eat-2* is expressed in most of pharyngeal muscles

It was reported that *eat-2* is the postsynaptic nicotinic ACh receptor gene for MC motor neurons and localized to small puncta near the junction of pharyngeal muscles pm4 and pm5 (19). Removal of wild-type bilateral MC neurons by laser ablation accelerated growth on CeMM, but the growth rate was still much slower than that of *eat-2* mutant (18). It is indicated that in addition to MC neurons, there might be other neurons synapse to EAT-2, which may play a similar regulatory role like MC neurons. Thus, we suspected that *eat-2* might have wider expression locations than as reported (19).

In order to observe the naïve expression state of *eat-2*. We used CRISPR/Cas9-mediated genome knock-in editing to insert a yellow fluorescent protein (YFP) into the intracellular loop between the third and fourth transmembrane domains of *eat-2* genome between leucine 377 and leucine 378 (SI Appendix, Fig. S1), and this CRISPR/Cas9 YFP knock-in strain was named *eat-2(rg552)*. Through spinning disk confocal fluorescence microscopy, we found that *eat-2* was expressed in multiple pharyngeal muscles, including pm1, pm3, pm4, pm5, pm6 and pm8 (Fig. 2A-C). This suggests that except for pm4, which was previously reported to connect MC neurons (18, 22). Those *eat-2* expressed pharyngeal muscle cells, which connect with other pharyngeal cholinergic neurons, such as M1, M2, M4 and M5. Thus, we suppose that M1, M2, M4 and M5 pharyngeal cholinergic neurons might also play a role in regulating growth similar to MC neurons.

**Fig. 2.**
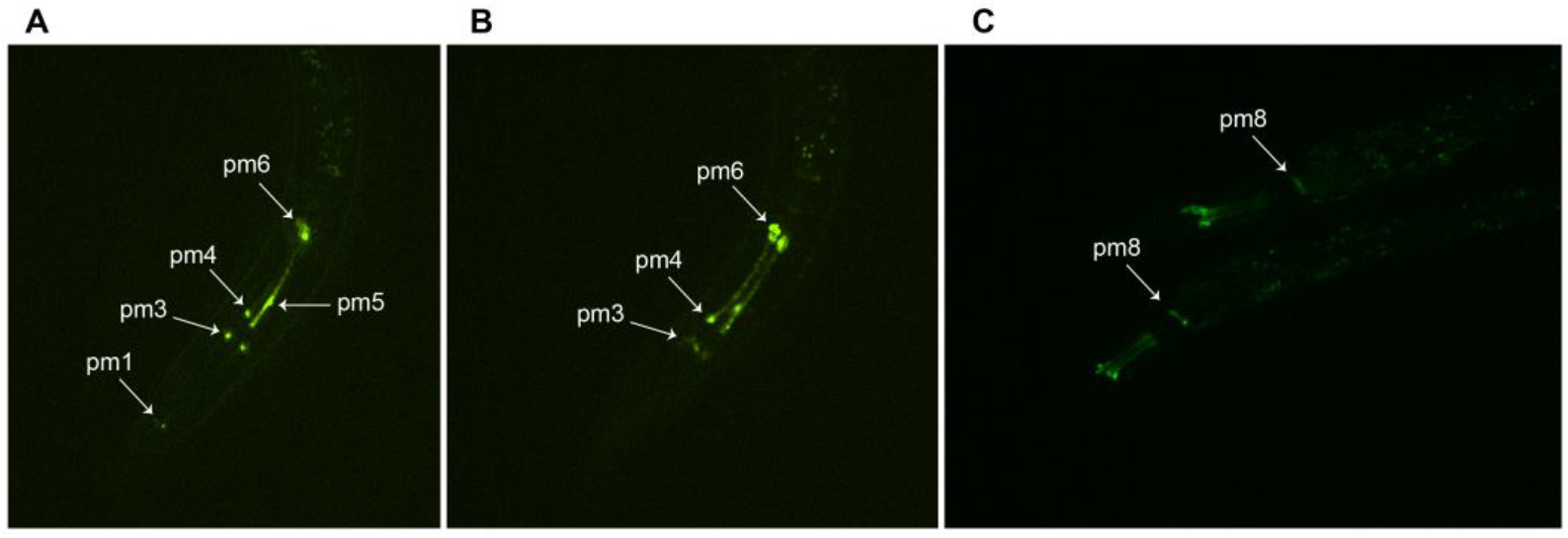
The expression of *eat-2* in pharyngeal muscle. (A-C) Spinning disk confocal microscope photographs. White arrows indicate expression of *eat-2* in pharyngeal muscles.

### Increased fatty acid anabolic metabolism in *eat-2* mutant on CeMM

To investigate how *eat-2* mutants promote worm growth downstream of pharyngeal cholinergic neuronal signaling. We performed transcriptome sequencing (RNA-seq) on N2 and *eat-2(ad1113)* mutant. Transcriptome sequencing results showed that after 2.5 hours of feeding on CeMM, the expression of 764 genes was significantly up-regulated and the expression of 1,673 genes was significantly down-regulated in *eat-2* mutant, compared with that of the wild type (SI Appendix, Fig. S2 A and B). Interestingly, KEGG pathway enrichment analysis of both up-regulated and down-regulated genes showed that fatty acid metabolism, fatty acid elongation, fatty acid degradation and biosynthesis of unsaturated fatty acids pathways were significantly enriched (Fig. 3A and B). This indicated that the growth difference between *eat-2* mutant and wild-type N2 might be associated with fatty acid-related metabolism.

**Fig. 3.**
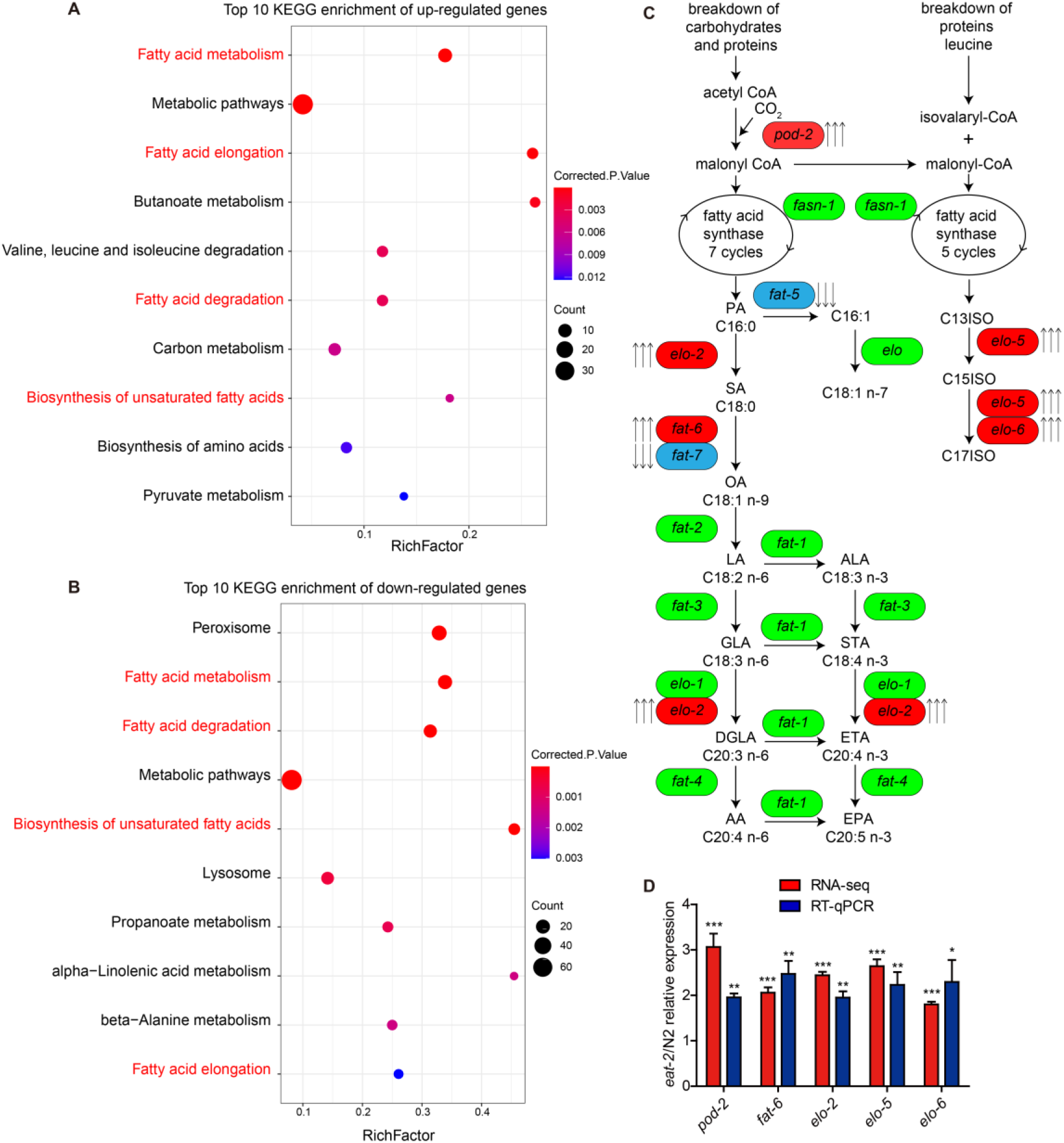
Expression changes of fatty acid metabolism related genes in *eat-2*. (A) The top 10 enriched KEGG pathways of significantly up-regulated genes in *eat-2* mutant. (B) The top 10 enriched KEGG pathways of significantly down-regulated genes in *eat-2* mutant. The significantly up-regulated and down-regulated genes were based on RNA-seq values. The rich factor is the ratio between the DEGs (differentially expressed genes) number and number of all genes in a certain pathway. Pathway of *de novo* fatty acid synthesis in *C. elegans*. This schematic diagram was adopted from reference (23). The three upward arrows indicate that the gene expression is significantly up-regulated in *eat-2* mutant, and the gene is marked in red. Three downward arrows indicate that the gene expression is significantly down-regulated in the *eat-2* mutant, and the gene is marked in blue. Genes marked in green indicate that their expression changes are not significant. The gene expression changes of *eat-2* mutant relative to wild-type N2 are based on RNA-seq results. ELO, elongase; C13iso, 11-methyldodecanoic acid; C15iso, 13-methyltetradecanoic acid; C17iso, 15-methylhexanoic acid; SA, stearic acid; OA, oleic acid; LA, linoleic acid; ALA, alpha linoleic acid; GLA, gamma linoleic acid; STA, stearidonic acid; DGLA, dihommo gamma linoleic acid; ETA, eicosatetraenoic acid; AA, arachidonic acid; EPA, eicosapentaenoic acid. (D) After feeding CeMM 2.5 h, fold difference in mRNA expression between wild-type N2 and *eat-2* mutant determined by RNA-seq and RT– qPCR. Three trials for RT-qPCR; data shown are mean ± s.e.m. \**P*<0.05, \*\**P*<0.01, \*\*\**P*<0.001. RT–qPCR results were statistical test by two-sided t test, RNA-seq results were analyzed by Wald test and adjusted for multiple testing using the Benjamini-Hochberg (BH) method.

Specially, further analysis found that, in *de novo* fatty acid synthesis process, except that *fat-5* and *fat-7* were down-regulated, the expression of several rate-limiting genes was significantly increased in *eat-2* mutant, including *pod-2*, *fat-6*, *elo-2*, *elo-5* and *elo-6* (Fig. 3C and D). Next, the differential expression of these five up-regulated genes was verified by reverse transcription quantitative real-time PCR (RT-qPCR). As expected, the results of RT-qPCR were consistent with RNA-seq, *pod-2*, *fat-6*, *elo-2*, *elo-5* and *elo-6* were indeed more abundantly expressed in *eat-2* mutant when fed with CeMM (Fig. 3D). Hence, we hypothesized that the enhanced fatty acid anabolism was responsible for the rapid growth of *eat-2* mutant on CeMM.

### Fatty acid synthesis genes are also increased in *tmc-1* mutant

We then wondered if other fast-growing mutant worms were also dependent on increased fatty acid synthesis. Following the method of studying *eat-2* mutant, RNA-seq for N2 and *tmc-1(rg1003)* mutant L1 larvae fed CeMM for 2.5 hours were performed. Compared with wild-type N2, the expression of 349 genes in *tmc-1* mutant was significantly increased, while the expression of 797 genes was significantly decreased (SI Appendix, Fig. S3 A and B). KEGG pathway enrichment analysis showed that fatty acid elongation and fatty acid metabolism were included in the top 10 up-regulated genes enriched pathways (SI Appendix, Fig. S4A), and fatty acid degradation was in the top 10 down-regulated genes enriched pathways (SI Appendix, Fig. S4B). Further analysis revealed that although *pod-2* and *fat-6* were not significantly upregulated (Data not shown), the expression level of *elo-2*, *elo-5* and *elo-6* were significantly elevated in *tmc-1* mutant (SI Appendix, Fig. S5). Subsequent RT-qPCR experiments also confirmed these trends (SI Appendix, Fig. S5). Collectively, we speculated that the fast-growth mutants accelerated the development of *C. elegans* on CeMM by increasing the fatty acid synthesis.

### Dietary fatty acid supplementation accelerates worm growth on CeMM

To test whether increased fatty acid synthesis actually accelerated the development of *C. elegans*. We added all the fatty acids (details are in Methods) on the *C. elegans de novo* fatty acid synthesis pathway (Fig. 3C) to CeMM one by one to detect their effects on worm growth. Surprisingly, when 1 mM of C17ISO (15-methyl-hexadecanoic acid), PA (palmitic acid) or SA (stearic acid) was added to CeMM, the growth rate of wild-type N2 worms was significantly increased (Fig. 4A, B and C). C17ISO can increase the adult rate from around 10% to 75% by 10 days post hatching, while either PA or SA could elevate the day 8 adult rate from about 2% to about 70% (Fig. 4A, B and C). Adding 0.2 mM acetyl CoA lithium salt, 1 mM C15ISO (13-methyl-tetradecanoic acid) and 1 mM OA (oleic acid) can also accelerate development, although not dramatically as C17ISO, PA and SA supplementation (Fig. 4D, E and F). Apart from this, the other fatty acids had no obvious effect on growth (SI Appendix, Fig. S6 A-H). PA and SA are substrate and product of *elo-2* elongated fatty acids, respectively (23). The synthesis of C17ISO requires action of fatty acid elongase *elo-5* and *elo-6* (23). Consistent with the significantly increased expression of *elo-2, elo-5* and *elo-6* genes in both *eat-2* and *tmc-1* fast-growing mutants (Fig. 3), our supplementation assays indicate that fatty acid synthesis metabolic reprogramming might regulate *C. elegans* development in the synthetic CeMM food environment.

**Fig. 4.**
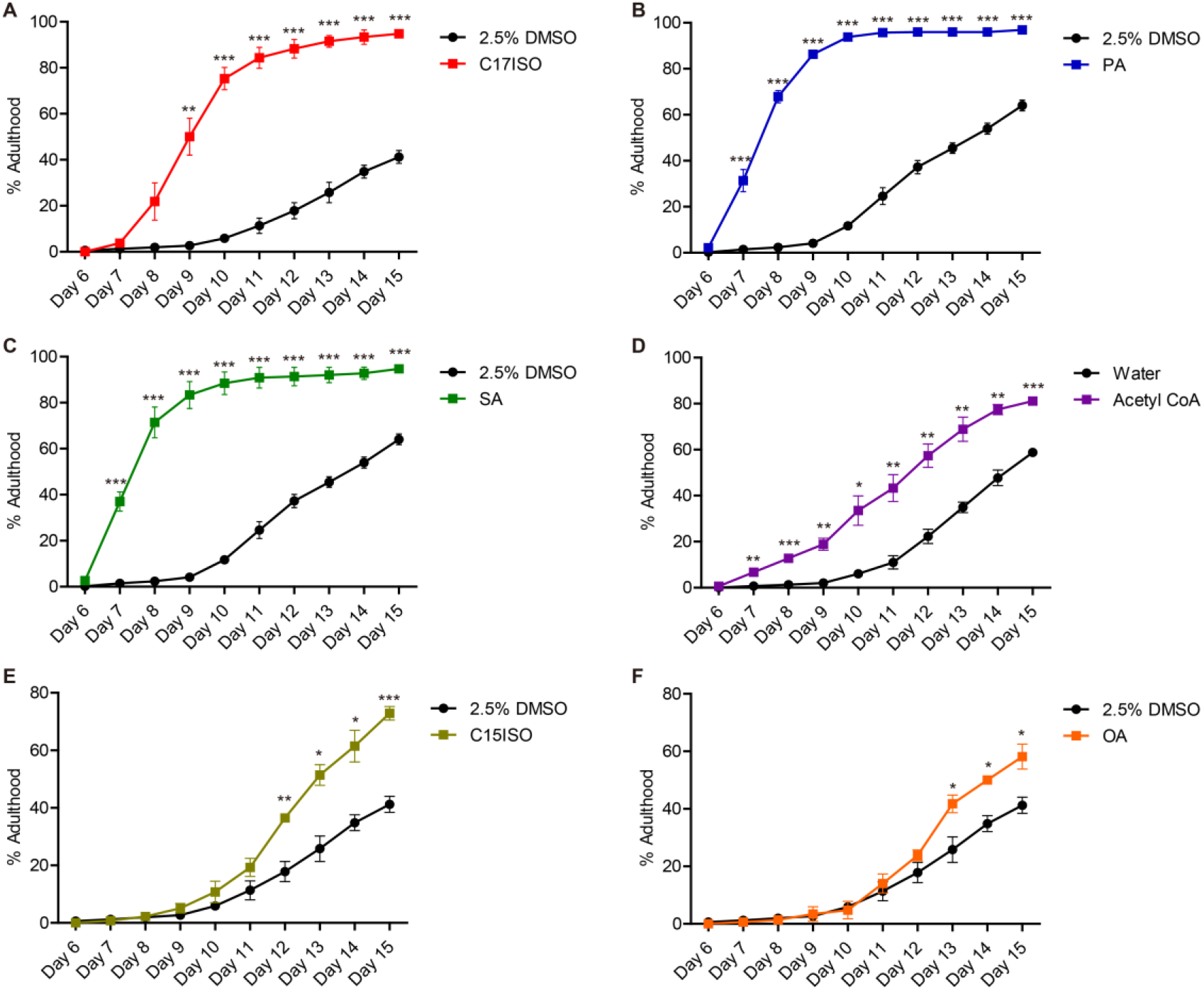
Supplementation of fatty acids to CeMM accelerated *C.elegans* growth. (A) After adding 1 mM C17ISO on the CeMM plates, the growth rate of wild-type N2 was accelerated. Control group 2.5% DMSO, three trials, n = 47, 53 and 53, respectively; C17ISO, three trials, n = 54, 48 and 54, respectively. (B) After adding 1 mM PA on the CeMM plates, the growth rate of wild-type N2 was greatly accelerated. Control group 2.5% DMSO, 8 trials, n = 44-55, 400 worms in total; PA, 8 trials, n = 46-58, 417 in total. (C) After adding 1 mM SA on the CeMM plates, the growth rate of wild-type N2 was greatly accelerated. Control group 2.5% DMSO, 8 trials, n = 44-55, 400 worms in total; SA, 8 trials, n = 44-58, 408 in total. (D) The growth rate of wild-type N2 nematodes after adding 0.2 mM acetyl CoA lithium salt. Control group water, three trials, n = 46, 48 and 56 worms, respectively; Acetyl CoA, three trials, n = 54, 55 and 55, respectively. (E) The growth rate of wild-type N2 nematodes after adding 1 mM C15ISO. Control group 2.5% DMSO, three trials, n = 47, 53 and 53, respectively; C15ISO, three trials, n = 49, 43 and 40, respectively. (F) The growth rate of wild-type N2 nematodes after adding 1 mM OA. Three trials were conducted in the two groups. Control group 2.5% DMSO, n = 47, 53 and 53, respectively; OA (C18:1 n-9), n = 52, 48 and 51, respectively. All data shown are mean ± s.e.m. \**P*<0.05, \*\**P*<0.01, \*\*\**P*<0.001; two-sided t test.

### *elo-6* regulates *eat-2* mutant development on CeMM

Since we found that *elo-2*, *elo-5* and *elo-6* expression were significantly increased in both *eat-2* and *tmc-1* fast-growing mutants (Fig. 3), together with the strong regulatory roles supported by our fatty acids supplementary assay (Fig. 4), we want to ask whether knocking them out actually attenuates the development of *eat-2* mutants. *elo-2*, *elo-5* and *elo-6* encode fatty acyl elongases and are homologs of *ELOVL3* (Elongation of very long chain fatty acids protein 3) and *ELOVL6* (Elongation of very long chain fatty acids protein 6) in human and mammal. *elo-2* knock-down by RNAi results in multiple phenotypic changes, such as slow growth, small body size, reproductive defects, and changes in rhythmic behavior (24). There was no published characterization on *elo-2* loss function of mutant, so we deleted *elo-2* in the *eat-2(ad1113)* mutant using CRISPR/Cas9 (SI Appendix, Fig. S7A). Due to the fact that *eat-2; elo-6* double mutant arrested at the L1 stage, we could not study the function of *elo-2* further. Previous studies have reported that *elo-5* plays a key role in the synthesis of monomethyl branched-chain fatty acids (mmBCFAs) C17ISO, and the absence of *elo-5* causes a severe L1 arrest phenotype (25). Consistent with this, the *eat-2; elo-5* double mutant we generated (SI Appendix, Fig. S7B) was also arrested in L1 larval stage, which prevented our further study. The suppression of *elo-6* activity by feeding double-stranded RNA (dsRNA) to wild-type animals did not cause obvious morphological or growth defects (25). We obtained three *elo-6* mutants with different alleles via CRISPR genome editing (SI Appendix, Fig. S7 C-E) in the background of *eat-2(ad1113)* mutant. Similar to *elo-6* knockdown as previously reported, the three alleles of *elo-6; eat-2(ad1113)* double mutants also behaved as superficially wild type, except that one of them, *eat-2(ad1113); elo-6(lzq11)* double mutant, was slightly shorter in body length. However, the growth rate of all the three *elo-6; eat-2(ad1113)* double mutants were significantly slower than *eat-2* single mutant on CeMM (Fig. 5A). On day 6 after hatching, about 57% of *eat-2(ad1113)* single mutant developed into adults, while only 4.3%, 8.3% and 27% of double mutant *eat-2(ad1113); elo-6(lzq19)*, *eat-2(ad1113); elo-6(lzq11)* and *eat-2(ad1113); elo-6(lzq12)* developed into adults, respectively (Fig. 5A).

**Fig. 5.**
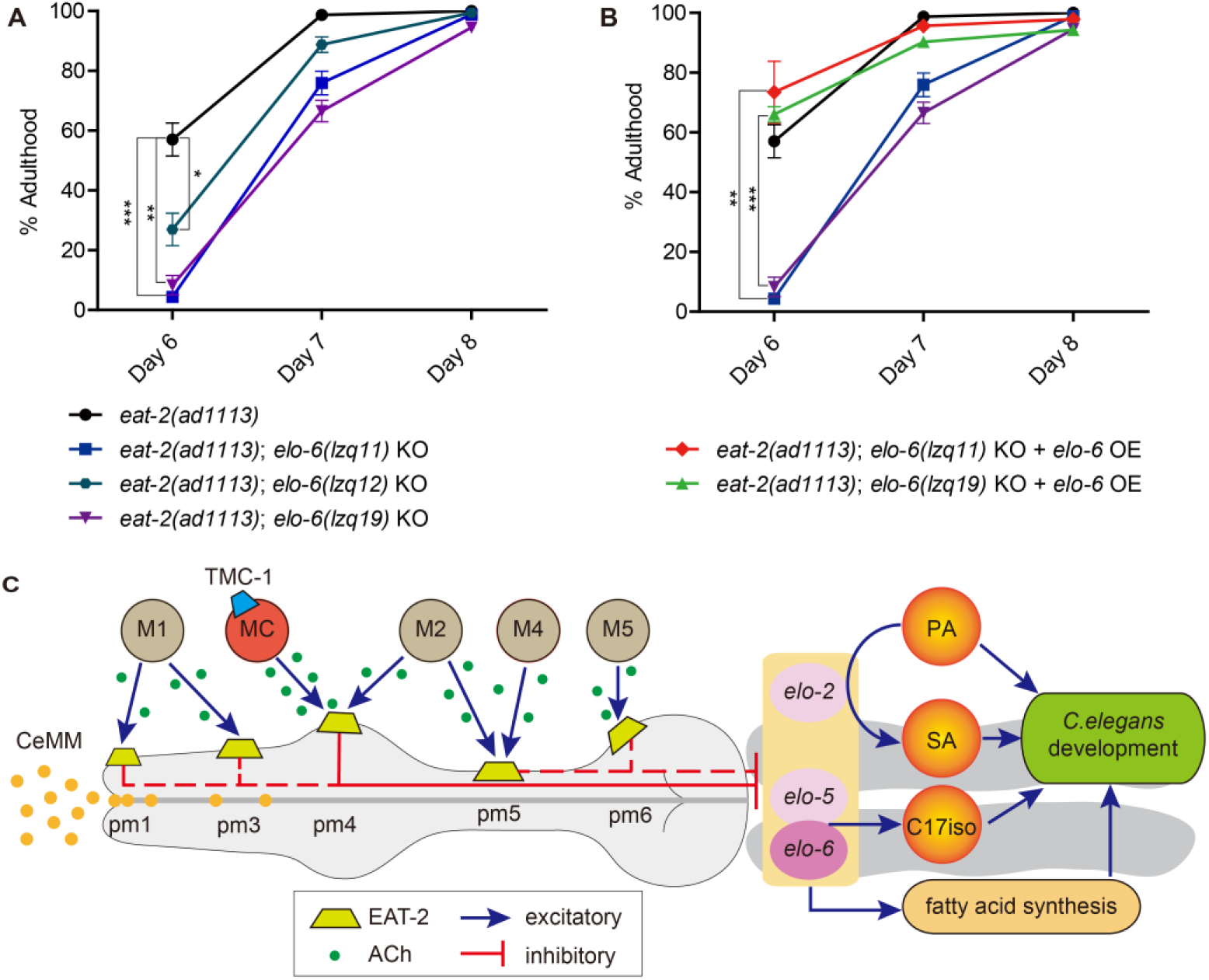
*elo-6* regulates *C. elegans* development. (A) Knockout of *elo-6* significantly slowed down the growth of *eat-2* mutant, *eat-2(ad1113)*, three trials, n = 53, 59 and 54 worms, respectively; *eat-2(ad1113); elo-6(lzq11)* KO, three trials, n = 59, 62 and 61, respectively; *eat-2(ad1113); eat-2(ad1113); elo-6(lzq19)* KO, three trials, n = 55, 55 and 58, respectively; *eat-2(ad1113); elo-6(lzq12)* KO, three trials, n = 59, 62 and 61, respectively. (B) *elo-6* overexpression could rescue the growth of *eat-2; elo-6* double mutants. *eat-2(ad1113)*, three trials, n = 53, 59 and 54 worms, respectively; *eat-2(ad1113); elo-6(lzq11)* KO, three trials, n = 59, 62 and 61, respectively; *eat-2(ad1113); elo-6(lzq11)* KO + *elo-6* OE, three trials, n = 55, 63 and 66, respectively; *eat-2(ad1113); elo-6(lzq19)* KO, three trials, n = 55, 55 and 58, respectively; *eat-2(ad1113); elo-6(lzq19)* KO + *elo-6* OE, three trials, n = 61, 63 and 72, respectively. All data shown are mean ± s.e.m. \**P*<0.05, \*\**P*<0.01, \*\*\**P*<0.001; two-sided t test. (C) Model depicts how acetylcholine signaling reprograms fatty acid synthesis and regulates *C. elegans* development. The gray color of M1, M2, M4 and M5 indicates that their regulatory function has not been confirmed. The red dotted line is the putative pathway of EAT-2 regulating growth and development. The neuromuscular connections shown in this figure were referred to WormAtlas and ref (26).

Then we want to know whether transgenic overexpression of *elo-6* can rescue the *eat-2; elo-6* double mutants’ slow growth on CeMM. We overexpressed the wild-type *elo-6* genomic DNA under the 1.4-kb *elo-6* promoter in the two of three double mutants: *eat-2(ad1113); elo-6(lzq19)* and *eat-2(ad1113); elo-6(lzq11)*, respectively. The results showed that overexpression of *elo-6* in both double mutants could significantly increase the growth rate to the level of *eat-2(ad1113)* single mutant (Fig. 5B). Collectively, the rapid growth of *eat-2* mutant depended on *elo-6*, which is in accordance to our C17ISO fatty acid supplementary test.

Together, our model (Fig. 5C) suggests that in wild-type N2 worms, MC and other pharyngeal cholinergic neurons such as M1, M2, M4 and M5, are excited on CeMM diets, and their acetylcholine neurotransmitter synapse to EAT-2 receptors located in multiple pharyngeal muscles; next the neuronal signals attenuates *C. elegans* development via downregulating fatty acid synthesis genes such as *elo-2*, *elo-5* and *elo-6*. In the absence of this negative regulation, fatty acid metabolism genes are significantly increased in both *eat-2* and *tmc-1* fast-growth mutants, compared to the wild-type animals. C17ISO, palmitic acid and stearic acid and several other fatty acids supplementations significantly accelerate *C. elegans* development, respectively. Essentially, *elo-6* is the key gene involved in the above fatty acid metabolism reprogramming and developmental regulating in *C. elegans*.

## Discussion

It was reported that impairment of cholinergic signaling increased food intake and lead to severe obesity, whereas enhanced cholinergic signaling reduced food consumption (4). Our results suggest that animals with null pharyngeal acetylcholine receptor *eat-2* grow much faster than wild-type on chemically defined synthetic CeMM, which is consistent with previous reports.

Our previous study demonstrated that TMC-1 has context-dependent functions, compared to N2 worms, *tmc-1* mutants grow much faster and have much higher male mating potency on CeMM food, while growth and male mating potency is similar between *tmc-1* mutants and wild-type N2 worms on OP50 food (18). Inhibition of growth by acetylcholine signaling seems to occur significantly only in CeMM nutritional environment, this suggests that genes like *eat-2* and *tmc-1* may regulate animal behavior in unfamiliar nutritional environments for animal adaptation in new ecological niche (18).

By knocking YFP into the genome *eat-2*, we found that *eat-2* was expressed in most of pharyngeal muscles. And these muscle cells are connected to multiple cholinergic neurons (Fig. 5C) (26). Therefore, we speculate that in addition to the previously reported MC-pm4 neuromuscular circuit (18), *eat-2* might regulate growth and development through multiple neuromuscular circuits. Further studies are needed to clarify the complete neural circuits that regulate diets sensation and development.

*C. elegans* can *de novo* synthesize fatty acids from acetyl-CoA, which is derived from breakdown of carbohydrates and proteins (23). However, detection by ^13^C isotope assay, it was found that only 7% of palmitic acid (16:0) in wild-type worms is derived from *de novo* synthesis, while the rest is directly absorbed from the bacteria food (27). More than 99% of *C. elegans* monomethyl branched-chain fatty acids (mmBCFAs) C17ISO was derived from de novo synthesis (27). Therefore, we hypothesized that wild-type worms could not obtain enough fatty acids from CeMM food, resulting in their developmental attenuation. Excitingly, our fatty acid supplementation experiments showed that dietary C17ISO, palmitic acid, stearic acid and several other fatty acids supplementation in CeMM significantly promoted the development and growth of wild-type N2 worms (Fig. 4).

C17ISO might act as a chemical/nutritional factor independent of the DAF-2/DAF-16 signaling pathway, and regulates post-embryonic development (28). C17ISO-lipid composition was reported to promote IP3 signaling, which in turn regulated membrane dynamics in the early embryo (29). In *C. elegans*, C17ISO was previously reported to be metabolized to 13-Methyltetradecanoic acid (C15ISO) that is used to synthesize sphingolipids (30). A mmBCFA-derived sphingolipid, d17iso-glucosylceramide (d17iso-GlcCer), is a critical metabolite in regulating postembryonic development via mTOR (31). Furthermore, C17ISO and d17iso-GlcCer were recently reported to mediate overall amino acid sensing through mTORC1 partly by controlling protein synthesis and ribosomal biogenesis (30). Our results showed that dietary supplementation of C17ISO can significantly promoted the development of *C. elegans* (Fig. 4A), and knocking out *elo-6*, a key enzyme involved in C17ISO synthesis, slowed down the rapid development of *C. elegans* (Fig. 5A). Given the broad existence of mmBCFA/sphingolipid in human diets and gut microbiota (32–34), together with the essential roles of sphingolipid in membrane homeostasis and related signalings such as IP3 and mTOR (29, 30, 35). And dysregulated metabolism of sphingolipids is linked to neurodegenerative processes in Parkinson’s disease, Alzheimer’s disease, amyotrophic lateral sclerosis (ALS) and Gaucher’s disease (36, 37), as well as obesity-related metabolic disorders such as diabetes, nonalcoholic fatty liver disease/steatohepatitis (NAFLD/NASH) and cardiovascular diseases (CVDs) (38). Further studies of the function of C17ISO might provide essential clues to understand how diets regulate human health and disease, and develop therapeutics to modulate ceramide levels to combat metabolic disease. Of note, we found that dietary supplementation of C15ISO also significantly promotes the development of wild-type *C. elegans*, but the growth acceleration rate is much lower compared to C17ISO supplementation on CeMM. To further elucidate why dietary supplementation of C15ISO and C17ISO behave differentially, is urgently needed.

We found that several fatty acyl elongases e.g., *elo-2*, *elo-5* and *elo-6*, were highly expressed in *eat-2* and *tmc-1* mutants (Fig. 3), which were fast-growing on CeMM. In particular, knock-out of *elo-6* in *eat-2* mutant slowed growth (Fig. 5A). Fatty acid elongases are conserved in animals, and *elo-2*, *elo-5* and *elo-6* are homolog of *ELOVL3* and *ELOVL6* in human and mammal. Related studies showed that *ELOVL3* plays a regulatory role in lipid recruitment in brown adipose tissue (39) and development of hair and skin function (40). *ELOVL6* not only regulates brown adipose tissue thermogenic capacity (41), but also affects insulin sensitivity (42, 43) and is associated with various diseases such as type 2 diabetes (42, 43), nonalcoholic steatohepatitis (44) and pulmonary fibrosis (45). Extensive studies suggest that fatty acid metabolism plays an important role in development, aging and disease (46–49). Our results suggested that diets sensed by the cholinergic neuronal circuits reprogrammed the fatty acid metabolism, which then regulated animal development and reproduction. Given most of the key genes and metabolites found in this study are conserved in both invertebrate and vertebrate animals, we believed that our results will provide important clinical clues to human health and diseases.

Interestingly, when a small amount of fatty acids is supplemented to the CeMM food, wild-type N2 worms significantly accelerate their development (Fig. 4), and the synchronization of development is enhanced, which will be more conducive to the application of drug screening using *C. elegans* and the improved CeMM food as a model platform. As *C. elegans* and CeMM platform have been employed for space biology studies, CeMM with fatty acid supplementation, developed in this report, will be widely accepted in the future space biology exploitations (50, 51).

Unexpected, after doing all the experiments, we discovered that we incorrectly used DL-lysine monohydrochloride (contains half of L-lysine monohydrochloride and half of D-lysine monohydrochloride) instead of L-lysine monohydrochloride. The decrease of L-lysine content and the introduction of D-lysine in CeMM medium caused the growth of all strains to be slowed down. Thus, we reformulated CeMM using L-lysine monohydrochloride, and the growth rates of wild-type N2 and *eat-2(ad1113)* mutant were increased to similar levels (SI Appendix, Fig. S8) as our previous study (18). Although the misuse of DL-lysine monohydrochloride in this study caused the growth of all strains to be slowed down, *eat-2* and *tmc-1* mutants still grew much faster than the wild-type animals on the above CeMM diet with DL-lysine, we believed that the mechanism of regulating growth and development in *C. elegans* remained unchanged. Given that decreasing of L-lysine and/or the introduction of D-lysine significant attenuates *C. elegans* development on CeMM, it is possible to further elucidate the functional roles of different types of lysine or the amount of lysine in *C. elegans* development.

In conclusion, this study demonstrates that fatty acid metabolism reprogramming regulated by the acetylcholine signaling modulates *C. elegans* development and growth in the chemically defined synthetic CeMM food environment, which could provide new insights on the molecular mechanisms underlying interactions among animal nutrition sensation, metabolism reprogramming and developmental regulation.

## Methods

### Strains and culture

Worms were cultivated at 20 °C on CeMM 1.7% agarose plates or OP50-seeded nematode growth medium (NGM) plates. Standard NGM is poured into disposable Petri plates and dried for 48 hours at room temperature before use. A complete list of strains is given in SI Appendix Table S1.

### CeMM preparation

The CeMM agarose plates were prepared as described by Zhang *et al*. (18). Dosages and manufacturers of all chemicals are in the SI Appendix Table S2. Hermaphrodites were age-assessed based on the developmental stage of vulva.

### Synchronization of *C. elegans*

Small-scale L1 synchronization was used for growth rate testing assays. Briefly, a drop of freshly prepared alkaline hypochlorite solution (50% bleach, 0.5 M NaOH) was put on one unseeded NGM plate, and then several gravid hermaphrodites from standard, well-fed culture stocks were transferred into this drop. After a few minutes, a second drop of alkaline hypochlorite solution was added to fully digest the adult body and bacteria. More gravid hermaphrodites can be bleached on other clean areas in the plate to obtain enough eggs. The plate was incubated at 20°C for ~18 hours for L1 offspring hatching.

Large-Scale L1 Synchronization was used for sample preparation of transcriptome and RT-qPCR. 90 mm NGM plates were seeded with 1 ml 15× concentrated OP50. The bacterial lawn was allowed to grown for 6 days at room temperature before transferring 35 synchronized L1s onto each seeded plate. Then, worms were cultured at 20°C and monitored daily until the plates were barely starved (i.e., there was a large supply of gravid hermaphrodites and minimal OP50 remaining on the plates, which can occur between 5 and 7 days). Worms were collected into several 50 ml conical bottom tubes with sterile water by washing off them from plates. The supernatant was removed after a few minutes of natural settling. Followed by additional washes until all bacteria were removed after which worms were collected as a ~400 μl pellet in several 15-ml conical tubes. Next, the worms were resuspended in 6 ml of low concentration alkaline hypochlorite solution (20% bleach, 0.5 M NaOH). The tubes were vortex for 5 s every 30 s to resuspend and facilitate dissolving of the adults. The dissolution of adults and release of eggs were monitored under a stereomicroscope. Sterile water was added to the alkaline hypochlorite solution to bring the volume to 12 ml when the adults was nearly completely dissolved (usually take 5-7 minutes). Eggs were pelleted via centrifugation at 1,300 × *g* for 1 minute after which supernatant was removed. The harvested eggs were washed 3 times with 12 ml sterile water, with vortex after every addition of sterile water. After the final wash, each tube of egg suspension with ~100 μl water was transferred to a 90 mm unseeded NGM plate using glass pipette. The plates were incubated for ~18 hours at 20°C to allow egg-hatching and checked the next day for synchronized L1s.

### RNA-seq and analysis

Synchronized L1s of different strains (N2, *eat-2(ad1113)* mutant or *tmc-1(rg1003)* mutant) were transferred to 90 mm CeMM agarose plates. Each plate contained ~25 μl L1 pellet which was suspended in ~100 μl of sterile water. Worms were allowed to grow for 2.5 h at 20°C. Treated worms were collected with sterile water by washing off them from CeMM plates followed by centrifugation at 1,150 × *g* for 1 minute. After the supernatant was removed, worm pellet samples were frozen in liquid nitrogen and then stored at - 80°C refrigerator. A biological duplicate includes ~50 μl pellet from two 90 mm CeMM plates. Triplicates were performed for each group of RNA-seq.

Total RNA was extracted using Trizol reagent kit (Invitrogen, Carlsbad, CA, USA) according to the manufacturer’s protocol. RNA quality was assessed on an Agilent 2100 Bioanalyzer (Agilent Technologies, Palo Alto, CA, USA) and checked using RNase free agarose gel electrophoresis. After total RNA was extracted, the mRNA was enriched by Oligo(dT) beads. Then the enriched mRNA was fragmented into short fragments using fragmentation buffer and reverse transcripted into cDNA with random primers. Second-strand cDNA were synthesized by DNA polymerase I, RNase H, dNTP and buffer. Then the cDNA fragments were purified with QiaQuick PCR extraction kit (Qiagen, Venlo, The Netherlands), end repaired, poly(A) added, and ligated to Illumina sequencing adapters. The ligation products were size selected by agarose gel electrophoresis, PCR amplified, and sequenced using Illumina HiSeq2500 by Gene Denovo Biotechnology Co. (Guangzhou, China). The raw reads were deposited into the NCBI Sequence Read Archive (SRA) database (Accession Number: PRJNA659845 for N2 and *eat-2(ad1113)* transcriptome; PRJNA659877 for N2 and *tmc-1(rg1003)* transcriptome).

Clean reads were obtained by removing adapters or low quality bases (reads with quality score ≤ 20) and ribosome RNA (rRNA) using short reads alignment tool Bowtie2 (52). An index of the reference genome was built, and paired-end clean reads were mapped to the reference genome by TopHat2 (53), respectively. Gene abundances of each group were quantified by software RSEM (54). The gene expression level was normalized by using FPKM (Fragments Per Kilobase of transcript per Million mapped reads) method.

To evaluate the reliability of experimental results as well as operational stability, the correlation analysis was performed by R. Principal component analysis (PCA) was performed with R package gmodels (http://www.rproject.org/) in this experience. To identify differentially expressed genes (DEGs) across samples, the DESeq2 software (55) was used. The genes with a fold change ≥ 1.5 and a false discovery rate (FDR) < 0.05 in a comparison were identified as significant DEGs. KEGG (Kyoto Encyclopedia of Genes and Genomes) pathway enrichment analysis of significant DEGs was completed by KOBAS (KEGG Orthology Based Annotation System) (56), the statistical test method was hypergeometric test / Fisher’s exact test, and corrected *P* value was calculated by Benjamini and Hochberg method.

### Real-time quantitative PCR (qPCR)

500 ng RNA per sample was used for reverse transcription using a cDNA synthesis kit (Toyobo, FSQ-301). Real-time PCR was performed on ABI QuantStudio 6 Flex system using SYBR Green (Toyobo, QPK-201). The internal reference gene was *act-1* in determining the gene expression difference between *eat-2(ad1113)* mutant and N2. And *mdh-1* was used as reference gene in detecting difference between *tmc-1(rg1003)* mutant and N2. Primers used for real-time PCR were shown in SI Appendix Table S3. All quantitative RT–qPCR reactions were performed at least in triplicate.

### Dietary supplementation

FAs C13ISO, C15ISO, C17ISO, C17anteISO, C16:0, C16:1 n7, C18:1 n7, C18:2 n6, C18:3 n3, C18:3 n6, C18:4 n3, C20:3 n6, C20:4 n3, C20:4 n6, C20:5 n3 (Larodan) and C18:0, C18:1 n9 (Aladdin), were prepared as 40 mM stocks in DMSO. The stock solution was mixed with sterile water in a 1:40 ratio. 100 μl of suspension was spread on the surface of one 3.5 mm CeMM plate, then the plate was allowed to dry in a 20°C dark incubator for one day before use. Acetyl-CoA lithium salt and malonyl-CoA lithium salt (Sigma) were prepared as a 10 mM stock in water. The stock solution was mixed with sterile water in a 1:40 ratio. Besides, STA (C18:4 n3) and ETA (C20:4 n3) stock solutions were dissolved in ethanol and diluted to 1 mm working concentration with sterile water before use. Similarly, 100 μl of the solution was spread on the surface of one CeMM plate, which was then dried in a 20°C dar k incubator for one day. The solvent, DMSO, water or ethanol, was used in control experiments.

### Molecular biology

The 4.6-kb *eat-2* genomic CDS was amplified with 5’-aatggacgatctagcATGACCTTGAAAATCGCATTT-3’ and 5’-aagagagaaaccgggTTATTCAATATCAACAATCGGACTA-3’ primers from N2 genomic DNA. The plasmid pMM1, which contained the *att* B Gateway destination cassette A, SL2 trans-splice site and red fluorescent protein, was PCR-linearized with 5’ - CCCGGTTTCTCTCTTTCTCTG-3’ and 5’-GCTAGATCGTCCATTCCGACA-3’ primers. Because the PCR primers that amplify the *eat-2* CDS sequence contain 15 bases of homology with the ends of the linearized vector, the *eat-2* CDS fragment was inserted between the linearized ends of pMM1, in a region after the Gateway destination cassette A, but before the SL2-red fluorescent protein site, using the In-Fusion HD EcoDry Cloning Kit (Clontech), to make the plasmid pXWC13.

The 4-kb upstream promoter fragment of *eat-2* gene was amplified from N2 genomic DNA. The sequences of the primers used were shown below: Attb1eat-2F1: 5’-ggggacaagtttgtacaaaaaagcaggctCAAATCGGCAAACCGGCAAATACA-3’, and Attb2eat-2R1: 5’- ggggaccactttgtacaagaaagctgggtCATGTAATAACTATCGCAATGTTTCG-3’. The primers contained Gateway *att*B sites, which allowed the 4-kb PCR products to be recombined, using BP clonase (Invitrogen), into the Gateway entry vector pDG15, to generate pLZ54. Then pLZ54 was recombined with pXWC13 using LR clonase (Invitrogen) to make the plasmid pXWC14.

The primer pair 5’-aatggacgatctagcATGCCACAGGGAGAAGTCTCATT-3’ / 5’-aagagagaaaccgggTTATTCAATTTTCTTTTCAGTCTTCTTC-3’ was used to PCR amplify the 1.8-kb *elo-6* genomic CDS. As described above, by using In-Fusion HD EcoDry Cloning Kit (Clontech), the *elo-6* CDS fragment was allowed to ligate into pMM1 to generate pXWC32.

The 1.4-kb upstream promoter region of *elo-6* (57) was amplified with primers with attB sites: 5’- ggggacaagtttgtacaaaaaagcaggctGGCGATTGTTGATTGTTGGTTTC-3’ / 5’- ggggaccactttgtacaagaaagctgggtTTTTACCTGCAATTTTAAACTTAAAAAAAG-3’. The amplified product was recombined into Gateway entry vector pDG15 through a BP recombination reaction (Invitrogen), then plasmid pXWC29 was generated. Finally, by using LR clonase (Invitrogen), pXWC29 was recombined with pXWC32 to obtain the expression plasmid pXWC36.

### CRISPR/Cas9-mediated genome editing

CRISPR/Cas9-mediated genome editing was carried out as described (58–60). Single-guide RNA (sgRNA) targeting sequences were predicted using online tool CHOPCHOP (https://chopchop.cbu.uib.no/) and individually cloned into the pDD162 vector (Addgene) by single site mutagenesis. sgRNAs used for generating *elo-2*, *elo-5* and *elo-6* mutant worms were as follows: *elo-2* sgRNA targeting sequence, 5’-GATGCATCAACTGGATTCTG-3’; *elo-5* sgRNA targeting sequence, 5’-GAATCTTTGAGATAACCCAA-3’; *elo-6* sgRNA1 targeting sequence, 5’-GGGAGAAGTCTCATTCTTTG-3’; *elo-6* sgRNA2 targeting sequence, 5’-GTGGAGAAGTCCAAATGTAG-3’. Molecular details of *elo-2*, *elo-5* and *elo-6* mutant alleles are shown in SI Appendix Figure S6.

The *eat-2* repair template, used for CRISPR/Cas9-mediated recombination, was generated by PCR amplifying from genomic DNA. A 1337 bp fragment containing the third and fourth transmembrane domains of *eat-2* genome between leucine 377 and leucine 378 was generated using the primers Attb1eat-2F3: 5’- ggggacaagtttgtacaaaaaagcaggctACTTCCTATCCGTGATGGTATTCC-3’ and Attb2eat-2R4: 5’- ggggaccactttgtacaagaaagctgggtTGAAACTTTACCAGTTTACTCGGT-3’. The fragment was then inserted into pDONR221 (Invitrogen) using BP clonase to generate pLZ72. We amplified YFP from a stock YFP-containing plasmid (pGW322) using the primers PGW322if-F1: 5’-GATGGCACGAAGTTGATGAGTAAAGGAGAAGAACTT-3’ and PGW322if-R1: 5’-TTGCTGGTTTTCAAGTTTGTATAGTTCATCCATGCCAT-3’. Using In-Fusion HD-cloning, YFP was translationally fused to *eat-2* genome between leucine 377 and leucine 378 in the plasmid pLZ72, to make the plasmid pLZ73.

To generate the *eat-2* CRISPR/Cas9 guide RNA plasmid pLZ90, the 20 bp guide RNA sequence to the *eat-2* (5’-GAGAAGAACGATGAAGAAGC-3’) was added to the CRISPR/Cas9 guide RNA/ enzyme plasmid pDD162 using PCR and the primers *eat-2* sgRNA F4: 5’-phosphate AGAAGAACGATGAAGAAGCGTTTTAGAGCTAGAAATAGCAAGT-3’ and sgRNA(universal)REV: 5’-CAAGACATCTCGCAATAGG-3’.

For genome editing in *eat-2*, the R324W mutation of *eat-2(ad1113)* was reconstructed in *eat-2(lzq10)*, the CRISPR sgRNA targeting sequence was: 5’-GCACCCAAAGACTCATCGGA-3’. The single strand DNA oligonucleotide repair donor used for this experiment was: 5’-CTCGATTTGCGCAAGTCTCATCATCGTCAACATTTTCTTCtGGCAtCCtAAaACaCAcAGaATGGGCGACTGGGTGAGCAATTTTGAAAATTTCTACAAA-3’. Lowercase letters represent different nucleotides from the genome, some of the sites in donor oligonucleotide were synonymously mutated to avoid Cas9-mediated cleavage of repair templates.

The co-CRISPR marker used was *rol-6* (61). The sgRNA targeting sequence is: 5’-GTTTAAAATGCAACGCTCTG-3’. It can create the *rol-6(su1006)* dominant mutation together with donor oligonucleotide, 5’- TGTGGGTTGATATGGTTAAACTTGGAGCAGGAACCGCTTCCAACCGTGTGc GctGcCAACAATATGGAGGATATGGAGCCACTGGTGTTCAGCCACCAGCAC CAAC-3’. For all CRISPR–Cas9-mediated genome editing assays, Cas9 target sites in donor molecules were synonymously mutated to avoid Cas9-mediated cleavage of repair templates. sgRNA vector (50 ng/μl) and repair template (50 ng/μl, if needed) were co-injected with *rol-6* sgRNA vector (50 ng/μl) and donor oligonucleotide (50 ng/μl). The final concentration of DNA in the injection mix did not exceed 200 ng/μl. Genome editing was confirmed by DNA sequencing.

### Generation of transgenic lines

All transgenic strains were generated by microinjection of the respective plasmid, and at least two independent lines of each transformation were used for experiments. Intestinal fluorescence expression plasmid pBL66 (P*gtl-1*::CFP) was added into the injection system as a co-transgenic marker. The detailed information of the transgenic strains is shown in SI Appendix Table S4.

### Statistical analysis

For graphs with error bars or statistical significance, detailed information of statistics and reproducibility are shown in the corresponding figure legends. Statistical analyses in this study were conducted using GraphPad Prism 5.

## Acknowledgement

We thank members of the L.Z laboratory for helpful discussions. Thanks also go to Profs. Jianhai Xiang, Fuhua Li and Baozhong Liu’ helps for suggestions, reagents and facilities. The work was supported by the“Marine life breakthrough funding” of KEMBL, Chinese Academy of Sciences; the “Young Scientist Research Program” of Qingdao National Laboratory for Marine Science and Technology; the Marine S&T Fund of Shandong Province for Pilot National Laboratory for Marine Science and Technology (Qingdao) [No. 2018SDKJ0302-1]; “Talents from overseas Program, IOCAS” of the Chinese Academy of Sciences; “Qingdao Innovation Leadership Program” [Grant 16-8-3-19-zhc]; and Key deployment project of Centre for Ocean Mega-Research of Science, Chinese Academy of Sciences. Some nematode strains were provided by the Caenorhabditis Genetics Center, which is funded by NIH Office of Research Infrastructure Programs (P40 OD010440).

